# Copper(II) Gluconate Boosts the Anti-SARS-CoV-2 Effect of Disulfiram *In Vitro*

**DOI:** 10.1101/2021.09.17.460613

**Authors:** Luyan Xu, Wei Xu, Simeng Zhao, Suwen Zhao, Lu Lu, Bo-Lin Lin

## Abstract

Disulfiram is a 70-year-old anti-alcoholism drug, while copper(II) gluconate (Cu(Glu)_2_) is a commonly used food additive or copper supplement. Here we disclose that the combination of disulfiram and copper(II) gluconate drastically enhances the anti-SARS-CoV-2 activity at the cellular level as compared to disulfiram or copper(II) gluconate alone. A 1:1 mixture of disulfiram and copper(II) gluconate shows an EC_50_ value of 154 nM against SARS-CoV-2 at the cellular level, much lower than the 17.45 μM reported for disulfiram alone. Notably, previous clinical trials have shown that a combination of 250 mg disulfiram (0.843 mmol) and 8 mg copper(II) gluconate (0.0176 mmol) oral capsules per day is well tolerated (NCT03034135, NCT00742911). A preliminary mechanism is proposed to rationalize the observed promotional effect.

---

We observed that the inhibitory effect of disulfiram (DSF) or copper(II) gluconate alone on SARS-COV-2 at the cellular level was 67.59% and 66.98% at 10 μM, respectively, while the inhibitory effect of a fresh 1:1 mixture of disulfiram and copper(II) gluconate was over 99% at 10 μM (**Table 1**). Further measurements showed that the EC_50_ of the 1:1 combination of disulfiram and copper(II) gluconate against SARS-COV-2 at the cellular level was 154 nM (**Figure 1**), significantly lower than that of disulfiram alone (17.45 μM).^1^

**Table 1.**
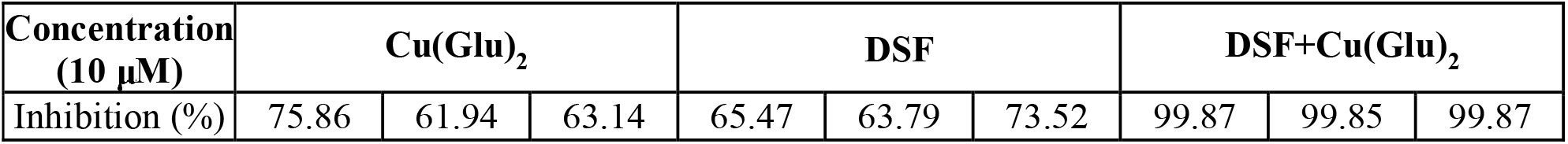
Inhibitory ratio of various chemicals by single-point inhibition assay at 10 μM onto SARS-COV-2 at the cellular level, determined by qRT-PCR analysis. n = 3 biological replicates.

| Concentration<br>(10 μM) | Cu(Glu)<sub>2</sub> | | | DSF | | | DSF+Cu(Glu)<sub>2</sub> | | |
| --- | --- | --- | --- | --- | --- | --- | --- | --- | --- |
| Inhibition (%) | 75.86 | 61.94 | 63.14 | 65.47 | 63.79 | 73.52 | 99.87 | 99.85 | 99.87 |

**Table 2.** EC_50_ values of various approved drugs against SARS-CoV-2 at the cellular level.

| Approved Drug | EC<sub>50</sub> (μM)<br>(Vero) | DOI |
| --- | --- | --- |
| **DSF+Cu(Glu)<sub>2</sub>** | **0.154** | **This work** |
| Boceprevir | 15.57 | 10.1038/s41467-020-18233-x |
| Disulfiram | ~10 | 10.1038/s41586-020-2223-y |
| Clofazimine | 0.31 | 10.1038/s41586-020-2577-1 |
| Atazanavir | 2.0 | 10.1021/acsptsci.0c00074 |
| Nelfinavir | 1.13 |  |
| Boceprevir | 1.9 |  |
| **Remdesivir** | **0.72** | 10.1126/science.abg5827 |
| Desloratadine | 0.7 |  |
| Flupentixol | 0.76 |  |
| Ttrimipramine | 1.5 |  |
| Lapatinib | 1.6 |  |
| Benztropine | 1.8 |  |
| Azelastine | 2.4 |  |
| Masitinib | 3.2 |  |

**Figure 1.**
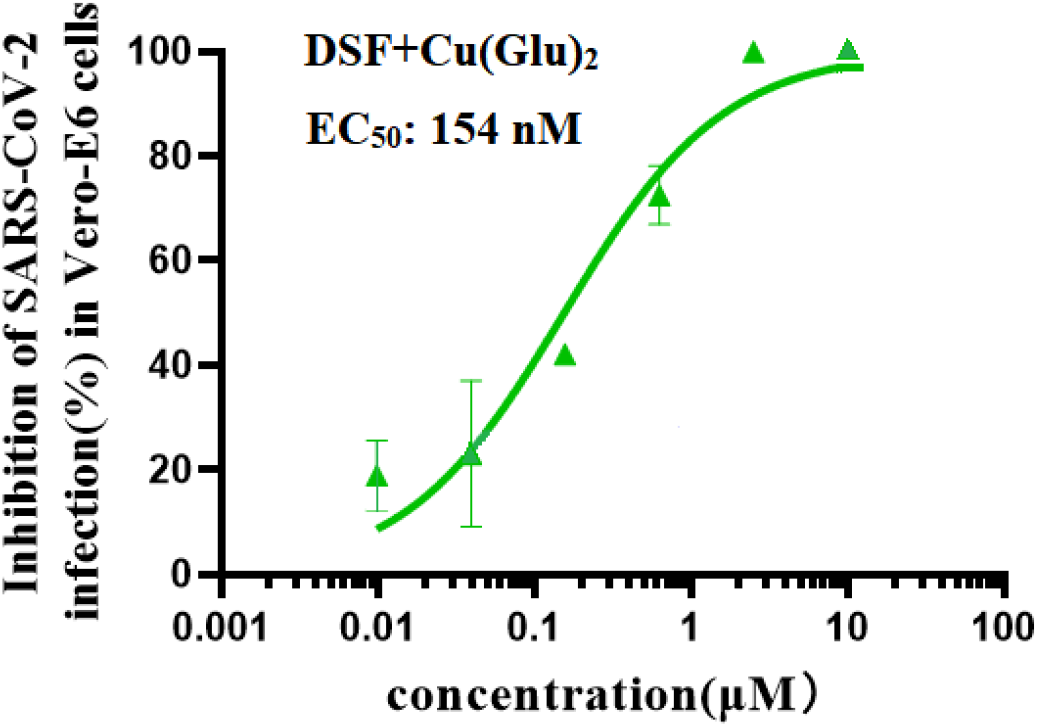
Dose-response curve for DSF+Cu(Glu)_2_, determined by qRT-PCR analysis, All data are shown as mean ± s.e.m., n = 3 biological replicates.

As shown in Figure 1, the mixture of disulfiram and copper(II) gluconate had much improved inhibitory activity of against SARS-CoV-2 infection with an EC_50_ of 154 nM (**Figure 1**). The results showed that viral RNA levels of the mixture of disulfiram and copper(II) gluconate were significantly lower than that of the non-mixture groups (disulfiram or copper(II) gluconate). Notably, previous clinical trials have shown that a combination of 250 mg disulfiram (0.843 mmol) and 8 mg copper(II) gluconate (0.0176 mmol) oral capsules per day is well tolerated (NCT03034135, NCT00742911). ^2, 3^

UV-vis spectroscopy revealed that the appearance and growth of an absorption peak at 433 nm from the 1:1 mixture of disulfiram and copper(II) gluconate over time, indicating the gradual formation of diethyldithiocarbamic acid cupric salt, Cu(DDC)_2_ (**Figure 2**). Dynamical light scattering spectroscopy showed that the resultant Cu(DDC)_2_ formed nanoparticles with the sizes of several hundred nanometers at the early stage (**Figure 3**). The particles slowly became larger than 1 micron and eventually precipitated out of the solution.

**Figure 2.**
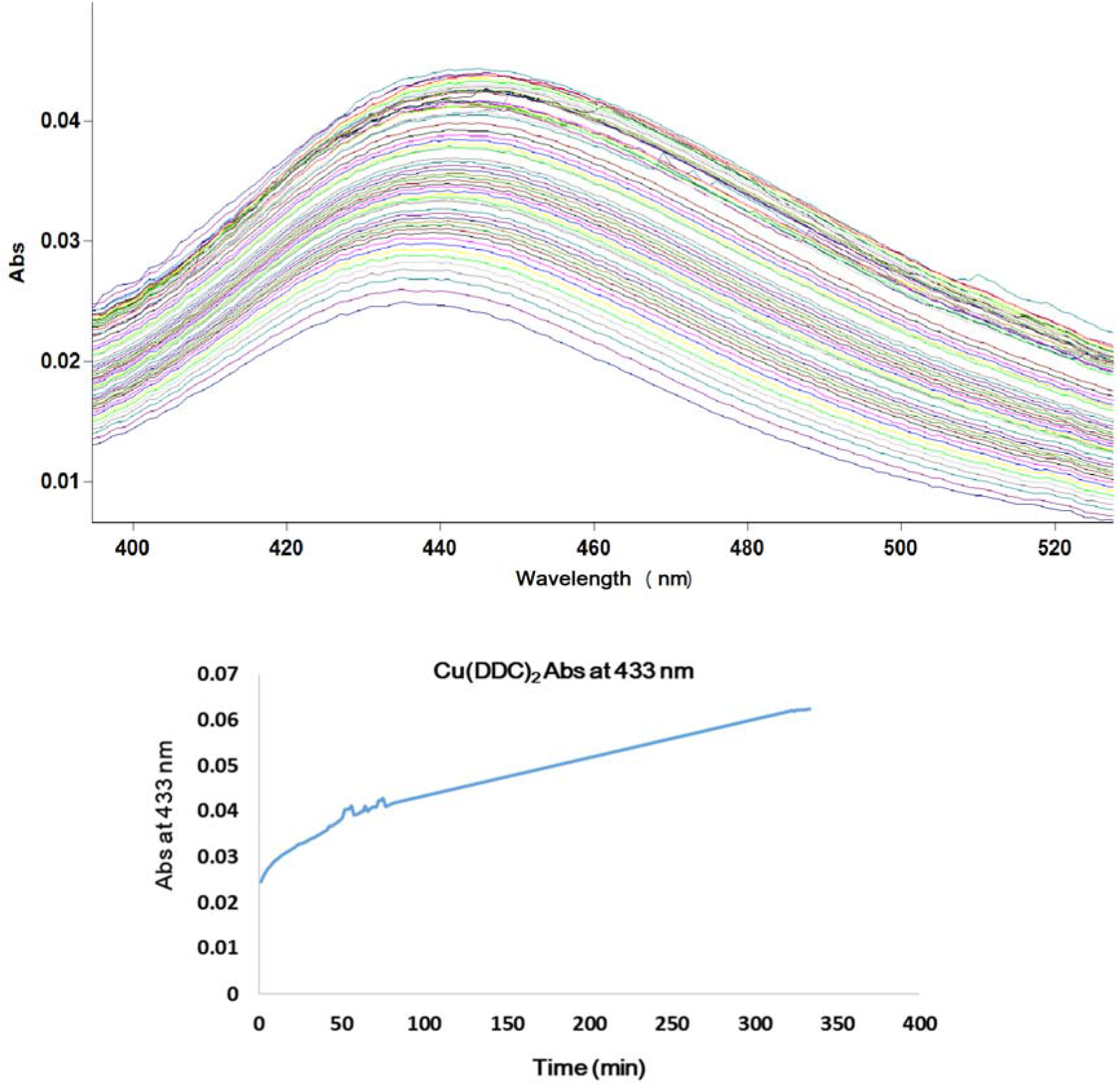
The appearance and growth of an absorption peak at 433 nm from the 1:1 mixture of disulfiram and copper(II) gluconate over time.

**Figure 3.**
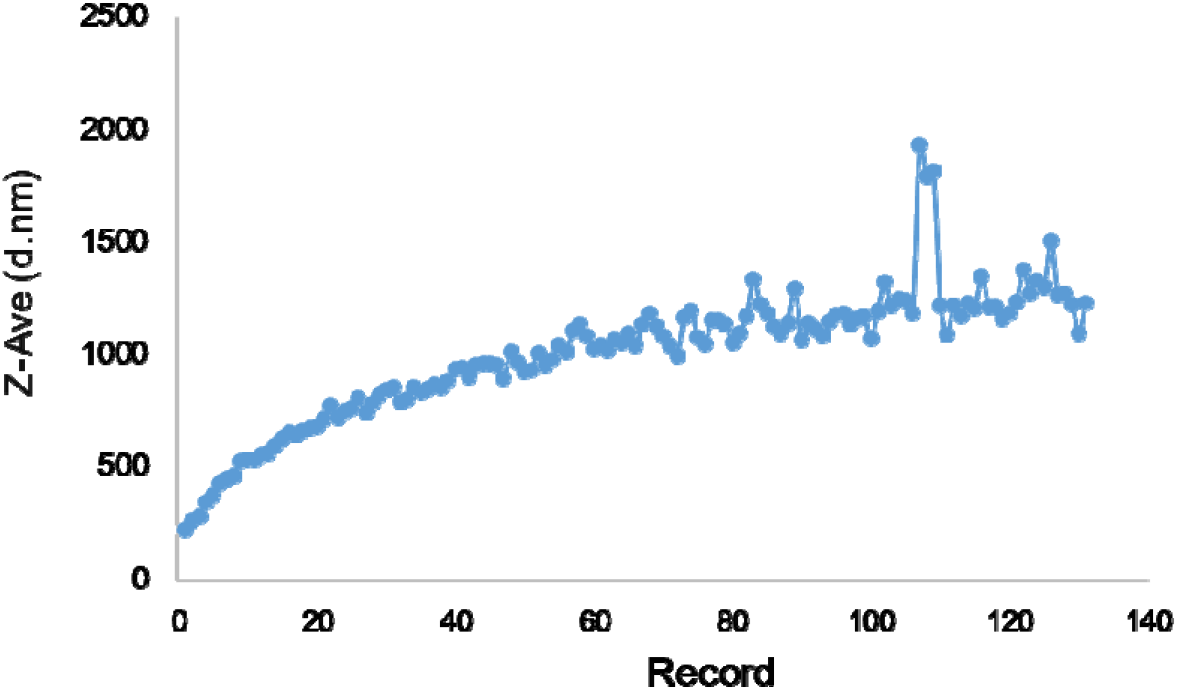
The growth of Cu(DDC)_2_ particles from the 1:1 mixture of disulfiram and copper(II) gluconate over time.

Notably, a comparison of some approved drugs shows that the combination of disulfiram and Cu(Glu)_2_ has a relatively low EC_50_ value against SARS-CoV-2 at the cellular level. However, further experiments, especially carefully designed clinical trials, are necessary before any conclusion can be draw on whether such combination can be used to treat COVID-19 or not.

Finally, we postulate that the boosting effect of Cu onto the anti-SARS-CoV-2 activity of disulfiram at the cellular level might be attributed to the *in situ* formation of Cu(DDC)_2_, which has been shown to possess an extraordinary ability to selectively oxidize zinc thiolate in the presence of thiols. Since zinc finger domains are present in several important SARS-CoV-2 enzymes crucial for the virus proliferation (*e.g.* RdRp),^4, 5^ Cu(DDC)_2_ may impair the function of these enzymes to inhibit SARS-CoV-2 proliferation via selective oxidation of the zinc thiolate sites of zinc finger domains even in the presence of a high concentration of glutathione.^6^

## Methods

### Anti-viral activity assays

A clinical isolate of SARS-CoV-2(SARS-CoV-2 / SH01 / human / 2020 / CHN, GenBank MT121215). The virus was purified by phagocytosis, reproduced in Vero-E6 cells, and stored in −80 ◻ for future use. All tests involving virus infection are strictly conducted at a biosafety level-3. Gradient-diluted drugs were mixed with SARS-CoV-2 (100TCID50) virus (the virus was diluted with serum-free DMEM medium) with the same volume, and incubated at 37 °C for 1 hour. In the 96-well plates (vero-E6 cells were 1×10^4^/ well), the supernatant was removed, 100 μL of the above drug/virus mixture was added, and the cells were incubated at 37°C for 1 hour. At the end of incubation, 100 μL of DMEM+2% FBS was added per well. After being placed in a cell incubator for further culture for 48 hours, the cell supernatant was collected for subsequent detection. Viral RNA was extracted from the cell supernatant using TRIzol LS reagent (Invitrogen) according to the manufacturer’s instructions. One-step PrimeScript™ RT Reagent Kit (Takara, Japan, Cat.#RR064A) Kit were used for quantitative real-time PCR. The reaction procedure of RT-PCR was: Reverse transcription: 95 ◻ 10s, 42 ◻ 5min; PCR reaction: (95 ◻ 5s, 56 ◻ 30s, 72 ◻ 30s)*40 cycles. The detection was carried out with a BIO-RAD quantitative fluorescence PCR instrument. The primers were: SARS-COV-2-N-F: GGGGAACTTCTCCTGCTAGAAT, SARS-CoV-2-N-R: CAGACATTTTGCTCTC AAGCTG, SARS-CoV-2-N-probe: 5’-FAM-TTGCTGCTGCTTGACAGATT-TAMRA-3’.

## Acknowledgements

We thank Dr. Qian Wang of Core Facility of Microbiology and Parasitology of Fudan University. We are very grateful to Dr. Di Qu, Xia Cai and Shan Su from Biosafety Level 3 Laboratory in Shanghai Medical College of Fudan University for their continuous support. This work was supported by the National Natural Science Foundation of China (No. U2032132).

## Competing interest

The authors declare no competing interests

## Notes

### Competing Interest Statement

The authors have declared no competing interest.

### Summary of Updates

Results of previous clinical trials on dosage tolerance of disulfiram plus copper gluconate are added into the summary and the text of the revised version.

